# Development of epigenetic clocks for New Zealand livestock

**DOI:** 10.1101/2021.06.30.450497

**Authors:** Alex Caulton, Ken G. Dodds, Kathryn M. McRae, Christine Couldrey, Steve Horvath, Shannon M. Clarke

## Abstract

Robust biological biomarkers of chronological age have been developed in humans and model mammalian species such as rats and mice using DNA methylation data. The concept of these so-called “epigenetic clocks” has emerged from a large body of literature describing the correlation between genome-wide methylation levels and age. Epigenetic clocks exploit this phenomenon and use small panels of differentially methylated cytosine (CpG) sites to make robust predictions of chronological age, independent of tissue type.

Here we present highly accurate livestock epigenetic clocks whereby we have used the custom mammalian methylation array “HorvathMammalMethyl40”^1^ to construct the first epigenetic clock for domesticated goat (*Capra hircus*), cattle (*Bos taurus*), Red (*Cervus elaphus*) and Wapiti deer (*Cervus canadensis*) and composite-breed sheep (*Ovis aries*). Additionally, we have constructed a New Zealand livestock ‘farm animal clock’ for all animals included in the study, which will enable robust predictions to be extended to various breeds. The farm animal clock shows similarly high accuracies to the individual species’ clocks (r>0.97), utilising only 217 CpG sites to estimate age (relative to the maximum lifespan of the species) with a single mathematical model.

We envision that the applications of this livestock clock could extend well beyond the scope of chronological age estimates. Many independent studies have demonstrated that a deviation between true age and clock derived molecular age is indicative of past and/or present health (including stress) status. There is, therefore, untapped potential to utilise livestock clocks in breeding programmes as a predictor for age-related, production traits.

## INTRODUCTION

In recent years, DNA methylation has been hailed as the most promising marker of biological age^3,4^. Despite the tissue-specific nature of age-associated physiological decline, cytosine methylation correlates strongly with age across virtually all tissue types. This unique characteristic can be exploited in the development of multivariate age estimators (pan-tissue epigenetic clocks) that can accurately predict the age of an individual independent of the tissue of origin of the DNA sample. For example, the first human pan-tissue clock, which is based on 353 age-related CpGs estimates chronological age with a correlation of 0.96 and median error of 3.6 years^5^. Since the development of this clock, epigenetic clocks have been successfully established for model species such as mice. More recently Lu et al. (2021) have constructed three universal mammalian epigenetic clocks from a large scale metanalysis of methylation data profiled using the HorvathMammalMethyl40 array^6^. The clocks show high accuracy (r>0.96) at predicting age across all 142 mammalian species and 57 tissue types assayed in the study, supporting the notion that a universally regulated mechanism underlies the aging process in mammalian species. This provides great potential for the use of an epigenetic clock for livestock breeding purposes. Establishing an accurate biomarker of age would be of use in farming systems where it is not possible to record birth date with certainty. In such circumstances, age cannot be reliably included in genetic prediction models, which results in reduced accuracy of genomic section estimates.

Perhaps of greater significance is that the estimates generated by this clock and its related counterparts are not only predictive of chronological age but also biological age, providing a mechanism to infer age-related pathologies and identify genetic, environmental or lifestyle factors that accelerate or slow biological aging^7^. For example, in humans, accelerated epigenetic ageing has been associated with a vast range of disorders from metabolic, infectious, and degenerative disease to frailty, traumatic stress and PTSD as well as lifestyle factors such as smoking and obesity^8-10^. These findings proffer the potential to utilise epigenetic clocks as a biomarker for health and age-related degeneration, a tool that could be deployed in the livestock sector to complement the current genetic framework for the selection of traits such as longevity (“stayability”)^11^. More specifically, early selection of animals with a “reduced biological age” could result in animals with improved longevity. Incorporation of the “biological age” in genetic evaluation to adjust an individual’s performance may enable better selection decisions. Alternatively, epigenetically younger animals may have a preference for retainment in the mob as a reflection of greater resilience. Furthermore, Lu et al. (2021) have shown that maximum lifespan, the genetic limit of longevity in an ideal environment, can also be predicted via methylation-based models^6^. Utilising the epigenetic clock as a biomarker in livestock breeding schemes would provide an objective predictor of stayability and health measures.

In New Zealand, the primary sector is dominated by the dairy cattle, beef cattle and sheep industries. In 2020 the number of beef cattle was 4 million, with 6.1 million dairy cattle, and 26.2 million sheep. The deer industry, comprised mainly of Red and Wapiti breeds, is well established for venison and velvet production, and total deer numbers are approaching 1 million. While goat farming has a lower profile, dairy goats are gaining popularity for their unique milk properties that are favourable for human infant formula, and currently, dairy goats number approximately 70,000. Breeding programmes are well established nationally across these key species, providing a strong foundation for the industry to capitalise on novel approaches to enhance the prevailing genetic merit predictions. Towards this goal, we have constructed an epigenetic clock for New Zealand farm animals to accurately estimate the chronological age of sheep, goat, deer and cattle (r=0.97). This work provides a tool to determine the biological age of livestock which will be particularly useful in farm systems and proffers potential as a molecular phenotype for stress resilience and age-related degeneration for breeding purposes.

## SAMPLES AND METHODS

### Deer samples

Although Red deer and Wapiti deer are classified as separate species, they are capable of interbreeding and produce fertile offspring. In the New Zealand deer industry Red and Wapiti are considered different breeds rather than different species, thus, to maintain consistency we will also refer to them as breeds and treat them as a single group for clock development^12^. Ear tissue punches (TSU samples) of deer were obtained from Focus Genetics, Napier, New Zealand. The animals were sampled from four different studs and were a mix of Red and Wapiti, while individuals in the dataset are not necessarily purebred, they are represented by their predominant breed. Ages ranged from 1 month old to 13 years and 8 months old. A total of 74 hinds and 22 stags were included in the dataset. Comprehensive birth date and sample date records enabled the age of the animal at the time of sampling to be determined to the nearest day.

### Cattle samples

TSU samples of cattle were obtained from LIC Ltd, Hamilton, New Zealand. The animals were sampled from the research dairy herd. The majority were Kiwicross breed (Jersey Friesian cross) with some full Jersey breed and some full Friesian breed included in the dataset. All 96 samples were female. Ages ranged from 1 month to 11 years and 1 month old, the age of the animal at the time of sampling was determined to the nearest day.

### Goat samples

TSU samples from one flock of Saanen breed dairy goats were obtained from New Zealand goat breeders, Judy and Barry Foote, Hikurangi, New Zealand. The dataset was made up of 72 does, 23 bucks and 1 intersex individual (visible male sex organs in addition to a milk-producing udder). Ages ranged from 2 weeks old to 8 years and 7 months of age, determined to the nearest fortnight.

### Sheep samples

TSU samples were obtained from two flocks of research sheep from AgResearch Ltd, NZ. The sheep were a composite breed resulting from the cross of Coopworth, Romney, Texel, Perendale and East Friesian. The age of the animals at the time of sampling ranged from 16 days to 7 years and 1 month old, determined to the nearest month. There were 73 ewes and 23 rams included in the dataset.

### Life-history traits

Life-history traits for the four species including maximum lifespan, average age at sexual maturity and average gestation length were sourced from the Animal Aging and Longevity Database (AnAge^13^, http://genomics.senescence.info/help.html#anage), (Table 1).

**Table 1:** Maximum life span and average life-history dates for deer, cattle, goat and sheep sourced from the AnAge database.

| **Species** | **Maximum Lifespan (years)** | **Age at sexual maturity (days)** | **Gestation length (days)** |
| --- | --- | --- | --- |
| **Deer** | 31.5 | 791 | 245 |
| **Cattle** | 20 | 548 | 277 |
| **Goat** | 20.8 | 545.5 | 155 |
| **Sheep** | 22.8 | 731 | 146 |

### Extraction and bisulphite conversion of DNA

TSUs were collected using ear punches and genomic DNA was extracted from tissue samples in 96-well plates using a high-salt method that has been shown to yield good quality, high molecular weight DNA^14^. Extraction and subsequent normalisation were performed with automated liquid handling robots.

Stock DNA was quantified using a Nanodrop8000 (Thermo Fisher Scientific Inc). The DNA was then diluted in autoclaved MilliQ water and normalised to approximately 80 ng/μl in a 96-well PCR plate. The normalised DNA plates were then quantified using an intercalating dye (PicoGreen; Invitrogen, USA) and an automated VICTOR3 fluorometer (Perkin Elmer, Inc., USA). Bisulphite conversion of 1ug input DNA was carried out with EZ DNA Methylation-Gold Kit (Zymo Research, USA) following the manufacturer’s instructions.

### Infinium array

A custom Illumina methylation array “HorvathMammalMethyl40” (Illumina, Inc. USA) was used to measure methylation at up to 37 thousand CpG sites, selected for their highly conserved flanking sequences across the Mammalia class. Array hybridisation and staining were performed with an automated liquid handling robot (Tecan Trading AG, Switzerland) and array scanning and imaging were executed on the Illumina Iscan platform.

### Data pre-processing

Raw .idat files were processed with the SeSAMe analysis suite to generate normalised beta values^15^. SeSAMe assesses the efficiency of probe hybridisation and extension using Infinium-I probe out-of-band measurements (the pOOBAH method) and effectively masks spurious methylation calls that result from incomplete probe hybridisation^15^. Normalised beta values are defined for each probe on a scale between zero (completely unmethylated) and one (completely methylated). SeSAMe has previously been implemented for normalisation of the Mammalian array and was shown to slightly outperform a similar R package minfi which implements the Normal-exponential convolution using out-of-band probes (NOOB) method of normalisation^1,16^.

Hierarchical clustering of the samples for each species was performed with the package WGCNA^17^ within R to establish if there was any underlying structure in the datasets that could be attributed to the known covariates including array batch effects or breed variation. Six sheep samples were excluded by quality control measures due to predicted sample contamination.

In line with the universal mammalian clock development, we constructed three epigenetic clocks for each species corresponding to three different transformations of age 1) **log-transformed chronological age**; 2) **-log(-log(RelativeAge))** and 3) **log-linear transformed age**^6^. Epigenetic age estimates of each clock were computed via the respective inverse transformation. The age transformations used to build clocks 1 to 3 incorporated three life-history trait measurements - gestational time (*GT*), age at sexual maturity (*ASM*), and maximum lifespan (*maxAge*), sourced from the AnAge database and measured in units of days (Table 1).

### Age transformations for clock construction

#### Log transformed chronological age for clock 1

For the log transformation of chronological age, an age offset of 2* gestational time was added to prevent logging zero based on the assumption that the epigenetic clock begins “ticking” from the moment of conception. Moreover, this avoids negative numbers in case that this clock was to be used on prenatal samples in future applications.

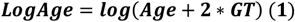

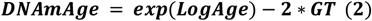

#### Loglog transformation of relative age for clock 2

Relative age, between 0 and 1, was defined using 2* gestational time, for similar reasons as clock 1, and maximum lifespan as follows:

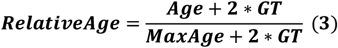

The loglog transformation of relative age then applied:

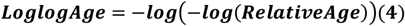

Clock 2 employs loglogAge and subsequently applies the inverse transformation to predict epigenetic age:

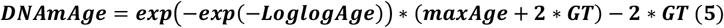

#### Log-linear transformation of age for clock 3

To define log-linear age, the average age at sexual maturity of the species is used. The transformation takes the logarithmic form when age is less than ASM and takes the linear form when age is greater than ASM and is continuously differentiable at ASM.

First, we calculate RelativeAdultAge as a ratio of age to ASM where an offset of 2* gestational time is added to ensure the RelativeAdultAge is always positive.

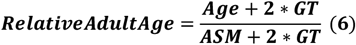

To model faster changes in methylation through development, a log-linear transformation on RelativeAdultAge was applied, samples were classified as young if their RelativeAdultAge was less than 1, hence Log-linear age is defined as:

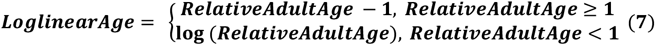

Clock 3 predicts *LoglinearAge* and applies the inverse transformation to estimate epigenetic age as follows.

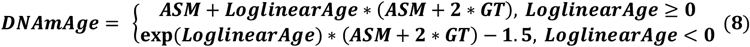

### Elastic net regression

To build epigenetic clocks for each species, an elastic net regression model was implemented using the R package glmnet^18^ within R v 3.626. Elastic net regression combines the regularisation of both ridge regression and LASSO (least absolute shrinkage and selection) regression making it suited to high dimensional datasets where the number of predictors is more than the number of observations and where predictors are likely to be highly correlated^18^. As part of the model, a subset of features (CpG methylation sites) are selected which cumulatively give the best predictor for a set of outcomes (age). The two main parameters that are employed in the elastic net model are; (1) the mixing parameter (α) which controls the shrinkage type between ridge (α = 0) and LASSO regression (α = 1) and; (2) the penalty parameter (λ) which controls the stringency of penalty (higher values of λ lead to coefficients closer or equal to zero)^19^.

The elastic net parameter α was set to 0.5 as the midpoint between both LASSO and Ridge regularization. The penalty parameter (λ) used was lambda.min calculated by cv.glmnet through a 10-fold internal cross-validation^20^. The mean error is calculated for each iteration of the cross-validation and then averaged over the ten partitions to determine the λ value that achieves the minimum mean error^18^.

The accuracy of the estimators was calculated using leave-one-out cross-validation (LOOCV) by executing the cv.glmnet() program on each set of n - 1 samples, where n is the number of samples. The predicted age of the omitted sample was calculated with the model built using data from the remaining samples.

### Farm animal clock

For the pan-species farm animal clock, we combined all data from all species and applied elastic net regression to the LoglogAge = -log(-log(RelativeAge)) as described in formulas (3), (4) and (5) to construct farm clock 2 and the loglinearAge transformation as described in formulas (6), (7) and (8) to construct farm clock 3. To assess the accuracy of the farm clocks, we used LOOCV and leave-one-species-out (LOSO) cross-validation. The LOSO cross-validation initially trained the model based on all-but-one species. The model was then tested on the omitted species to assess how well it generalized to the species that were not included in the training dataset.

### Assessing the model prediction performance

To validate the accuracy of the models, we determined the Pearson’s correlation and calculated the median absolute difference (median absolute error; MAE) between the epigenetic age estimates from LOOCV analysis and the observed age for all samples. The correlations and MAE were also computed for the LOSO-based approach for each species. The correlations were calculated on the transformed data (where the relationship was linear).

## RESULTS

### Structure of the raw data

To initially characterise any substructure in the datasets we performed hierarchical clustering using the full methylation array information for each sample. There were four distinct clusters based on species with moderate sub clustering based on sex, importantly there was no obvious sub clustering based on batch effects from samples run on the same chip array (Figure 1).

**Figure 1:**
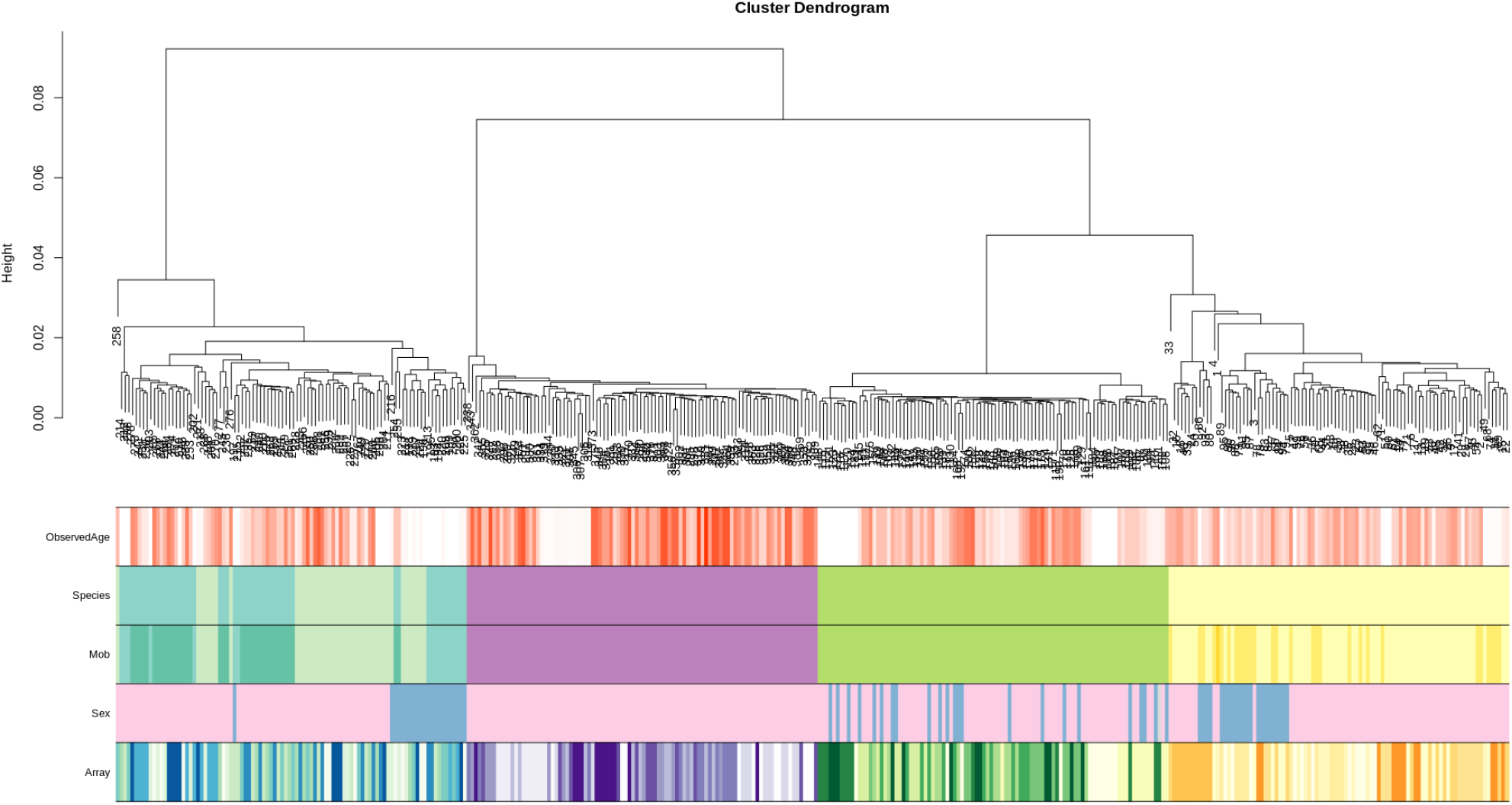
Hierarchical clustering of the complete methylation dataset for each sample (excluding sheep samples that failed QC). Species are coloured yellow for sheep related covariates, green for goat related covariates, purple for cattle related covariates and turquoise/blue for deer related covariates (note: the two shades of turquoise in the species panel are representative of Red and Wapiti breeds, respectively. Sex is coloured pink for female, blue for male and grey for hermaphrodite samples. The data forms four distinct clusters based on species, there is some sub clustering based on sex, there is no obvious sub clustering attributed to the array the sample was run on.

**Figure 2:**
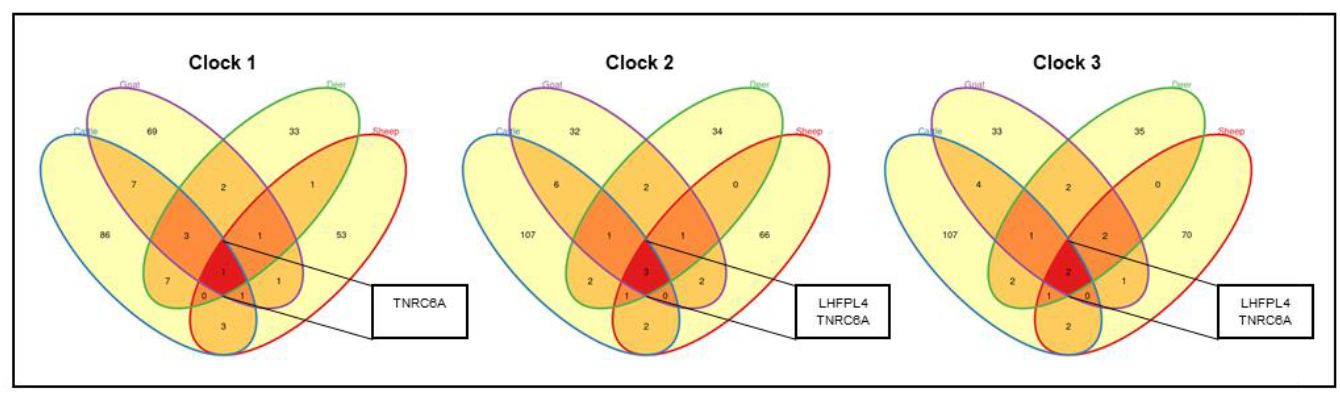
Venn diagrams depicting the overlap of CpG sites included in each of the individual species’ clocks for clock 1, clock 2 and clock 3. The CpG sites common across all four clocks lie within the genes TNRC6A and LHFPL4.

### Development of species-specific clocks

We constructed three epigenetic clocks per species based on three different transformations of age which differed slightly in their output and respective accuracies. The first epigenetic clock (clock 1) correlates epigenetic age with observed chronological age. An age offset of 2* gestational time is added to the observed age to account for circumstances when the clock may be applied to prenatal samples. Clock 2 was developed to allow for biologically meaningful comparisons to be made between the four species by accounting for their differing maximum lifespans. Clock 2 leverages both gestational time and maximum lifespan to determine age relative to maximum age on a scale between 0 and 1.

Clock 3 uses log-linear transformed age and accounts for greater changes in methylation during early life development. This clock leverages age at sexual maturity as a proxy for maximum lifespan as it correlates strongly with maximum lifespan on the log scale (Pearson correlation r=0.82, p=6×10^-183^ across all mammalian species in AnAge).^6^ Age at sexual maturity is often better characterised in animals compared to maximum lifespan, particularly in livestock that are typically not retained beyond their productive lifespan.

The number of CpG sites selected by elastic net regression to construct the models for each species ranged from 45 to 123 sites. On average 60% of the sites for each clock were negatively and 40% positively correlated with age, respectively. (Table S1)

### Predictive performance of the epigenetic clocks

All three clocks for each species were highly accurate (r>0.937) with an MAE of less than 7 months (Figure 3). Clock 1 consistently, albeit slightly, outperformed both clock 2 and clock 3, which can be expected given that clock 2 and 3 make assumptions regarding maximum lifespan and age at sexual maturity. The goat clock 1 was the most accurate (r=0.992) using only 86 CpG sites for predictions. The high accuracy of this clock is likely due to the controlled environment in which the goats were raised and their origination from a single, closed herd.

**figure 3:**
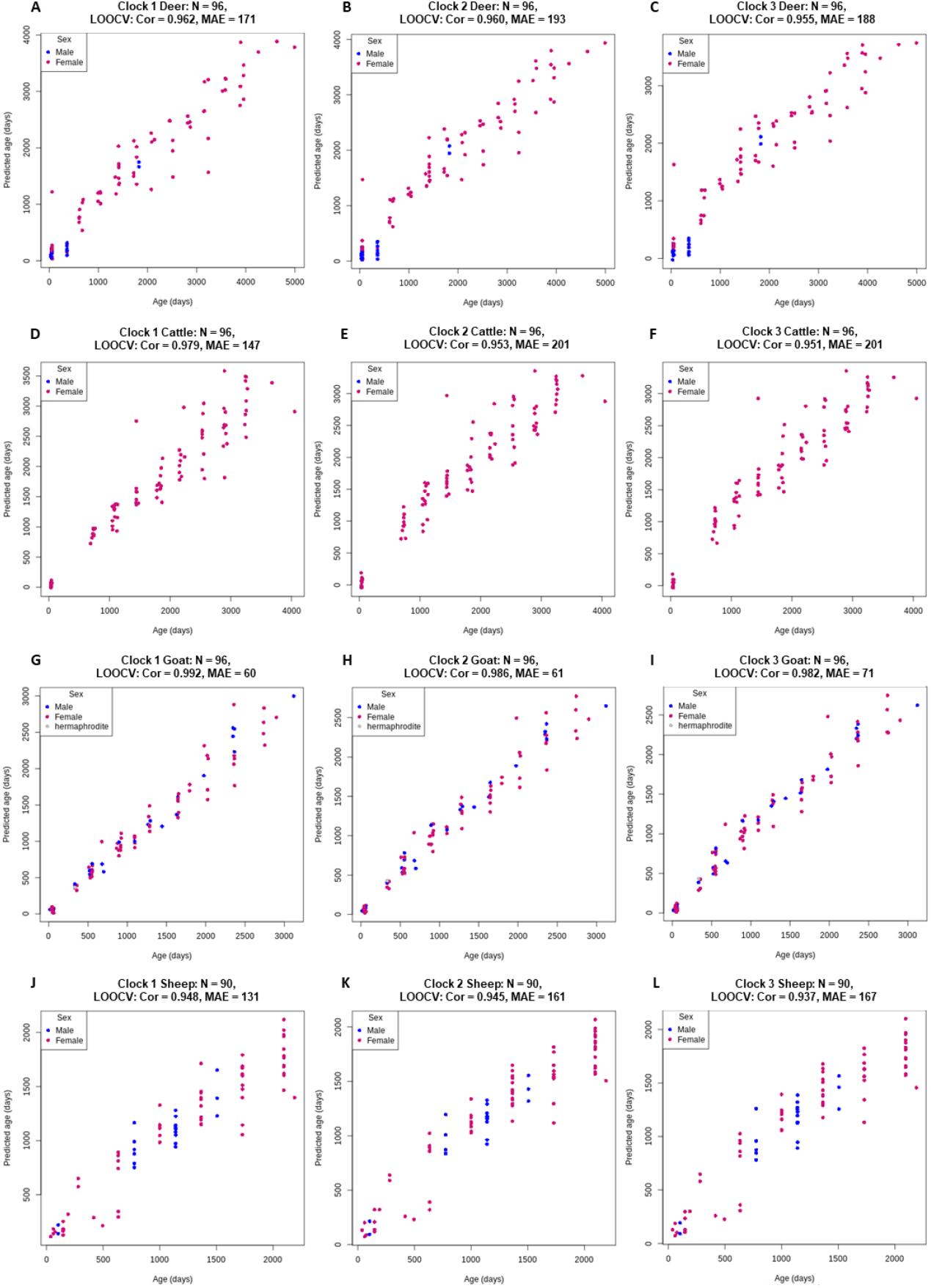
Back transformed chronological age (x-axis) versus back transformed epigenetic age estimated for clock 1 (log transformed chronological age; **A,D,G,J**), clock 2 (loglogRelativeAge; **B,E,H,K**) and clock 3 (log-linear transformed age; **C,F,I,L**). For all four species, deer (**A, B, C**), cattle, (**D,E,F**), goat (**G,H,I**) and sheep (**J,K,L**). The Pearson correlation coefficient estimates (Cor) and median absolute error (MAE) are reported for the age estimates via LOOCV.

To improve the robustness of the individual clocks we built a pan-species clock using the complete dataset of all four species. For the farm animal clock, we performed both a LOOCV and a LOSO cross-validation to assess its performance. For the LOSO approach, one species was omitted per iteration and the model built with the remaining three species before being tested on the omitted group. The performance of the farm animal clock 2 shows similarly high accuracy to the individual species clocks using 217 CpG sites with a median correlation of 0.97 for the LOOCV cross-validation and more notably, a median correlation of 0.94 from the LOSO analysis (Figure 4, Table S2).

**Figure 4:**
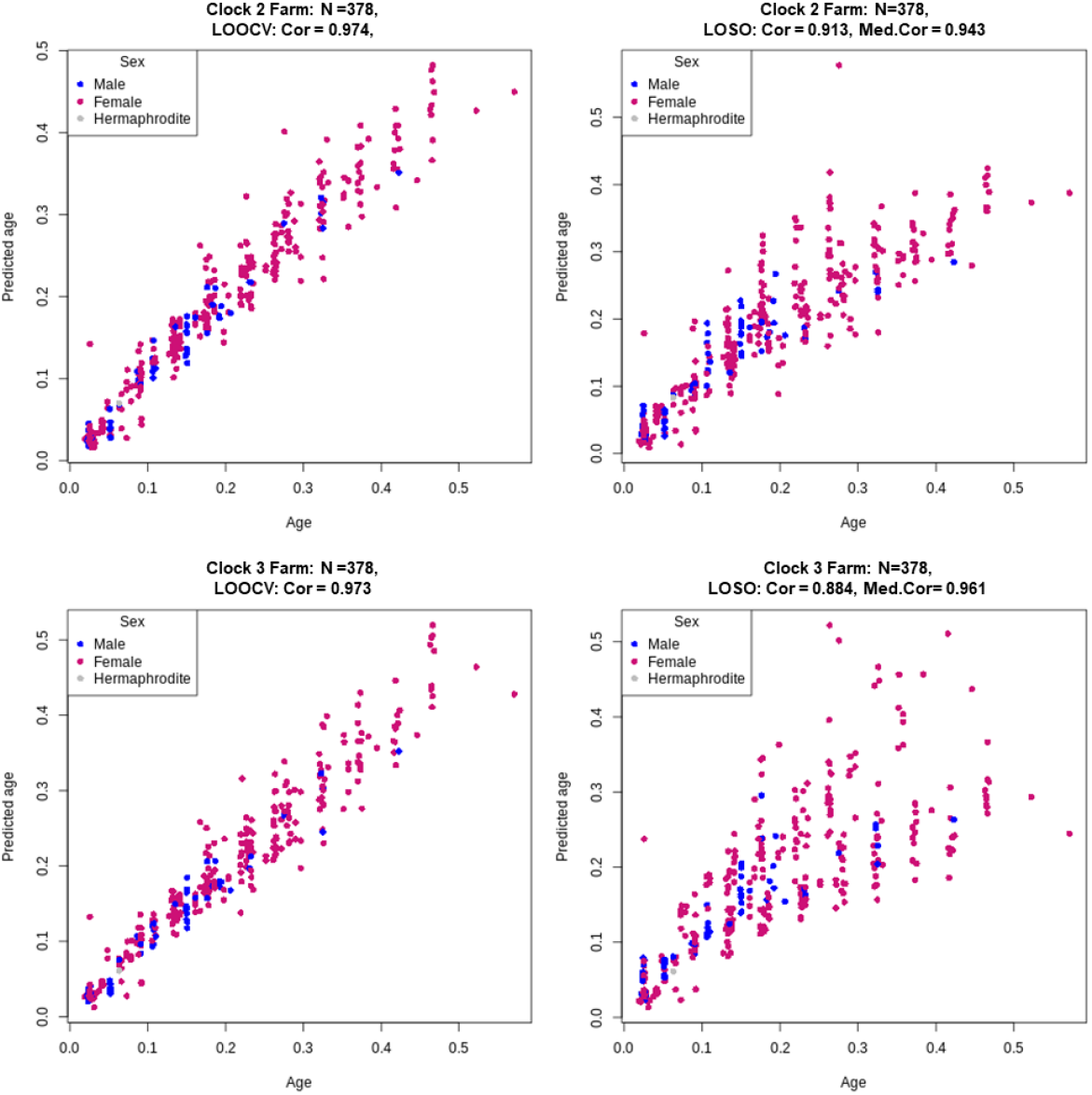
back-transformed relative age (x-axis) versus back-transformed relative epigenetic age estimated for Farm clock 2 (loglogRelativeAge; **A,B**) and Farm clock 3 (log-linear transformed age; **C,D**). Age estimates were predicted via both LOOCV (A,C) and LOSO (B,D) cross-validation. The Pearson correlation coefficient estimates (Cor) are reported for the age estimates via both LOOCV and LOSO cross-validation and the median correlation coefficients (Med.Cor) for each fold of the LOSO cross-validation are reported.

At first interpretation, the individual species clocks may appear to be highly biased towards the species used to construct the clock, resulting in inflated accuracy estimates. However, the remarkable predictive accuracy of LOSO analysis refutes this notion and demonstrates that the pan-species farm animal clock is highly robust and likely to perform well even in some species that are not incorporated in the training dataset and/or in different environments, for example in livestock originating from different countries.

### Overlap between CpG sites

There was very little overlap between the CpG sites selected for the clocks of each species. Only one CpG site from clock 1, three CpG sites from clock 2 and two CpG sites from clock 3 were shared across all four species, respectively; these were located in two functionally annotated genic regions, TNRC6A and LHFPL4 (Figure 2).

TNRC6A encodes a member of the trinucleotide repeat containing 6 protein family and was consistently demethylated with increasing age. The protein functions in post-transcriptional gene silencing through RNA interference (RNAi) and microRNA pathways^21^. Demethylation of this gene has previously been linked to ageing in similar epigenetic clock studies in prairie voles^22^ and primates^23^. The LHFPL4 gene is a member of the lipoma HMGIC fusion partner (LHFP) gene family, which is a subset of the superfamily of tetraspan transmembrane protein encoding genes. LHFPL4 plays a role in the regulation of inhibitory synapse formation and maintaining GABA receptors^24^. While there are no obvious links between its function and the ageing process, two cytosines located in exon 2 of in the LHFPL4 were the most predictive across all species in the development of the recent universal mammalian epigenetic clock and showed a correlation with age of >0.8 in 24 species, implying there is a true, as of yet unclear function of this gene and its role in the ageing process^6^. Further work by Lu et al., 2021 revealed that methylation of LHFPL4 cg12841266 was strongly correlated with both development (r=0.58 and P=8.9×10-11) and post-development stages (r=0.45 and P=2.3×10-76) across a range of tissue types. Importantly the observed changes in methylation levels of LHFPL4 were consistently observed to be associated with ageing in young animals as well as both middle-aged and old animals suggesting that this gene plays a role throughout the lifespan of mammals from development through to adulthood^6^.

## DISCUSSION

The premise of using biomarkers to predict longevity emerged from an understanding that chronological age is an imperfect surrogate of the ageing mechanism and a more informative measure is the decline in functional capability with age, that is, an individual’s biological age^2^.

In the livestock sector, the longevity of an animal is of significant economic importance and also represents a measure of animal welfare and sustainability within the sector^25^. Extending the productive lifespan of an animal increases the profitability of breeding schemes by (1) reducing the cost of replacement animals; (2) improving the average mob yields by increasing the proportion of animals in higher-producing age groups, for example a larger proportion of mature cows in dairy cattle schemes; (3) optimising land usage by reducing acreage required to rear replacement animals; and (4) decreasing involuntary culling (culling productive, profitable animals due to illness, injury, infertility) while increasing voluntary replacement (culling or selling animals that are healthy but not meeting productivity requirements)^26^.

Selection based on longevity is complex, as the true lifespan of an animal is only available at the end of its natural life while breeding decisions and culling occur earlier. Longevity or “staybiulty” is used to reflect the capability of an animal to remain in the mob over time, avoiding both natural attrition and culling. Identifying biomarkers that are predictive of an animal’s functional stayability would greatly enhance the current selection framework for this complex trait. Epigenetic clocks, which are predictive of age-related degeneration in mice and humans, could be an appropriate biomarker towards this goal.

This paper outlines the development of a novel epigenetic clock for New Zealand farm animals, providing a tool to accurately estimate the chronological age of sheep, goat, deer and cattle with high accuracy (r>0.97), with obvious application to additional livestock species. The results from this work echo the key findings of the universal mammalian epigenetic clock that was trained on a large-scale dataset across 128 mammalian species. Firstly, and most noteworthy, is the robustness of both the farm and universal clocks at predicting biological age across species that were not part of the training set, which reinforces the notion that a highly conserved and defined mechanism underlies biological ageing. Secondly, the *LHFPL4* gene which is highly predictive of age in both the universal mammlain clocks and the farm animal clocks, would appear to play a key role in the ageing process throughout an individual’s lifespan, despite its current functional annotation providing little connection to biological degeneration. The lack of overlap between the CpG sites selected for the clocks of each species can likely be explained by the LASSO element of elastic net regression whereby if there are grouped variables (highly correlated between each other) LASSO tends to select one variable from each group ignoring the others^27^. Therefore, if there are two or more correlated predictors one may be randomly selected for a specific clock while reduced to zero in a different clock.

We expect that the livestock clock will prove a useful tool for the livestock industry to make accurate predictions of biological age and envision the clock sites could be incorporated into a small-panel, targeted assay to reduce the costs associated with array-based DNA methylation profiling. This study provides a tool to determine the biological age of livestock with huge potential as a molecular phenotype for age-related degeneration and stayability traits for breeding purposes.

## ACKNOWLEDGEMENTS AND FUNDING

This work was primarily supported by the Ministry of Business Innovation and Employment, New Zealand award C10X1906 “Beyond the genome: Exploiting methylomes to accelerate adaptation to a changing environment” (Clarke S.) and AgResearch Ltd SSIF funding. The goat array-typing was supported by Paul G. Allen Frontiers Group (Horvath S.). Caulton A. was supported by a University of Otago Ph.D. scholarship. The funders had no role in study design, data collection and analysis, decision to publish, or preparation of the article. The Mammalian Methylation Consortium authored the Universal Mammalian clock article from which this study’s methods were based. The following people and corporations kindly supplied the tissue samples: Barry and Judy Foote supplied the goat samples, LIC supplied the cattle samples, Focus Genetics supplied the deer samples and AgResearch suppled the sheep samples.

## AUTHOR CONTRIBUTIONS

Caulton A. and Dodds K. G developed the clocks. Caulton A. carried out additional bioinformatics analyses. Horvath S. and his lab group generated the array data for the goat samples. Caulton A. generated the array data for cattle, deer and sheep. All authors helped with editing the article and data interpretation. Clarke S. conceived of the study.

## SUPPLEMENTARY DATA

**Table S1:**
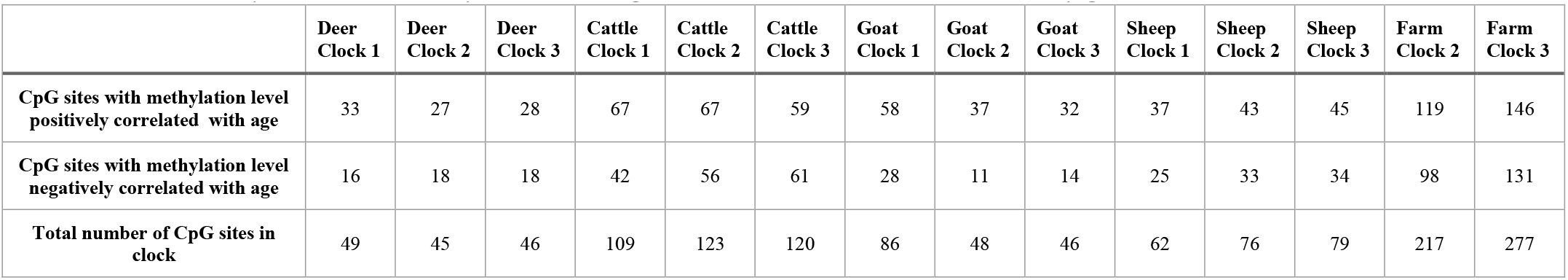
Number of CpG sites selected by elastic net regression for the construction of each epigenetic clock.

|  | Deer<br>Clock 1 | Deer<br>Clock 2 | Deer<br>Clock 3 | Cattle<br>Clock 1 | Cattle<br>Clock 2 | Cattle<br>Clock 3 | Goat<br>Clock 1 | Goat<br>Clock 2 | Goat<br>Clock 3 | Sheep<br>Clock 1 | Sheep<br>Clock 2 | Sheep<br>Clock 3 | Farm<br>Clock 2 | Farm<br>Clock 3 |
| --- | --- | --- | --- | --- | --- | --- | --- | --- | --- | --- | --- | --- | --- | --- |
| **CpG sites with methylation level positively correlated with age** | 33 | 27 | 28 | 67 | 67 | 59 | 58 | 37 | 32 | 37 | 43 | 45 | 119 | 146 |
| **CpG sites with methylation level negatively correlated with age** | 16 | 18 | 18 | 42 | 56 | 61 | 28 | 11 | 14 | 25 | 33 | 34 | 98 | 131 |
| **Total number of CpG sites in clock** | 49 | 45 | 46 | 109 | 123 | 120 | 86 | 48 | 46 | 62 | 76 | 79 | 217 | 277 |

**Table S2:** The Pearson correlation coefficients for the age estimates via LOSO cross-validation for each fold drop for clock 2 and clock 3. The data is trained on 3 species and tested on the omitted species. For each iteration we report the correlation between the transformed epigenetic age and the transformed age of the omitted species.

| Test species | LOSO CV correlation (Clock 2) | LOSO CV correlation (Clock 3) |
| --- | --- | --- |
| **Deer** | 0.93 | 0.84 |
| **Cattle** | 0.96 | 0.95 |
| **Goat** | 0.98 | 0.92 |
| **Sheep** | 0.92 | 0.89 |

